# Structural Basis for the transmembrane signaling and antidepressant-induced activation of the receptor tyrosine kinase TrkB

**DOI:** 10.1101/2023.06.06.543881

**Authors:** Erik F. Kot, Sergey A. Goncharuk, María Luisa Franco, Alexander S. Arseniev, Andrea Benito-Martínez, Mario Costa, Antonino Cattaneo, Marçal Vilar, Konstantin S. Mineev

**Author notes:** These authors jointly supervised the work. Current address: Goethe University Frankfurt, Frankfurt am Main, 60438, Germany (Konstantin S. Mineev). These authors contributed equally to the work (Erik F. Kot, Sergey A. Goncharuk, María Luisa Franco).

## Abstract

Neurotrophin receptors of the Trk family are involved in the regulation of brain development and neuroplasticity, and therefore can serve as targets for anti-cancer and stroke-recovery drugs, antidepressants, and many others. The structures of Trk protein domains in various states upon activation need to be elucidated to allow rational drug design. However, little is known about the conformations of the transmembrane and juxtamembrane domains of Trk receptors. In the present study, we employed NMR spectroscopy to solve the structure of the TrkB dimeric transmembrane domain in the lipid environment. We verified the structure using mutagenesis and confirmed that the conformation corresponds to the active state of the receptor. Subsequent study of TrkB interaction with the antidepressant drug fluoxetine, and the antipsychotic drug chlorpromazine, provided a clear self-consistent model, describing the mechanism by which fluoxetine activates the receptor by binding to its transmembrane domain.

## 1. Introduction

Trks are a family of high-affinity neurotrophin receptors belonging to receptor tyrosine kinases. Trks proteins regulate neuronal survival and differentiation^1,2^, and are involved in a variety of diseases, including cancers and neurodegenerative disorders, which makes them prospective drug targets^3,4^. The Human Trk family includes three members: TrkA, TrkB, and TrkC, each specific to its own neurotrophin molecule: NGF, BDNF, and NT3^5^. The structures and molecular mechanisms of Trk activation are mostly known. Extracellular domains of Trks have been studied separately and in complex with ligands by NMR and X-ray crystallography^6–10^, revealing a peculiar ligand-binding mode. These parts of Trks consist of five subdomains, and only the C-terminal Ig-like domain (d5) is involved in ligand binding, whereas the roles of the other four are still mostly unknown. Moreover, the ligands protrude from the d5 domains towards the cell membrane by approximately 2 nm and most likely interact with the disordered extracellular juxtamembrane regions ^11^. The latter was found to play a significant role in regulating Trk activity ^12–14^. The structures of the kinase domains of various Trks were also obtained by crystallography and revealed the fold typical for receptor tyrosine kinases^15^.

One of the most intriguing parts of Trk receptors is their transmembrane domains. Until recently, Trks were thought to be activated by ligand-induced dimerization. However, first, by cross-linking^16,17^ and later by single-molecule FRET approaches^18^, Trks were shown to exist as pre-formed dimers on the cell membrane, suggesting that dimerization itself does not induce receptor activity. This mechanism seems to be common for various kinds of receptor tyrosine kinases, and the TrkB receptor provides the highest population of pre-formed dimers among all the receptors tested ^19^. Altogether, this implies that there are two dimeric states of Trk transmembrane domains (TMDs), corresponding to the active and inactive forms of the receptor. The structure of TrkA TMD in one of these states, presumably inactive, was previously solved by NMR spectroscopy. The second state was characterized by cross-linking, functional assays and TOXRed assays^20^. Switching between the two conformations, induced by cross-linking in the juxtamembrane regions, has been directly observed in the most recent work^11^. However, the atomistic details of the TMD dimer, corresponding to the Trk active state, and the TMD conformations of other Trk members besides TrkA, as well as the TMD conformations of other Trk members besides TrkA, remain elusive.

The functional role of Trk TMDs goes beyond receptor homodimerization or switching between the two conformations. TMDs can mediate protein-protein interactions, as demonstrated by the example of TrkA and p75 receptors^21^. TrkA can modulate the activity of the p75 neurotrophin receptor, and the TMDs of both proteins were shown to play an essential role in this phenomenon. Finally, the most peculiar is the TrkB receptor, which has been shown to bind several antidepressant drugs belonging to different chemical classes and is involved in neuroplasticity in response to antidepressant therapies^22^. Surprisingly, according to recent studies^23^, the binding of antidepressants occurs in the transmembrane part of the receptor and is closely related to the protein-cholesterol interactions in cell membranes. Moreover, TrkB TM domain was recently also shown to be a high-affinity binding site for psychedelic drugs LSD and psilocin, the active metabolite of psilocybin, that have shown antidepressant potential in clinical trials^24^. This makes the TMD of TrkB an interesting drug target and highlights the need for atomistic structural data describing this region. In this regard, here we report the structural investigation of the TrkB transmembrane domain in lipid bicelles, accompanied by direct analysis of its interactions with the environment and several commercial antidepressant drugs employing NMR spectroscopy in solution.

## 2. Results

### 2.1. Spatial structure of TrkB transmembrane domain in the lipid environment

It is well established that the structures of helical transmembrane dimers may be affected by the choice of membrane mimetic, and in some cases, switching between active and inactive states may occur upon transfer from one environment to another^25–27^. Additionally, the effects of cholesterol and antidepressant drugs must be studied in the presence of a lipid bilayer to ensure the proper position of these molecules with respect to the TM domain. For this purpose, in the current work, we synthesized the transmembrane domain of human TrkB (TrkBtm, residues 423-466, UniProt ID Q16620) and reconstituted it in the classic lipid-containing environment of DMPC/DHPC bicelles as described^24^. Under these conditions one can observe two sets of cross-peaks in the HSQC spectra of TrkBtm (S_1_ and S_2_ below), with intensities dependent on the lipid-to-protein ratio (LPR) (Figure 1 A, B, S1). This behavior is typical for oligomerization; therefore, we applied water/bicelle dilution and ^1^H/^15^N cross-relaxation analyses^28^ to establish the order of the respective oligomers. As revealed by titration, oligomer S_2_ is the dimer of oligomer S_1_, and the bicelle hydrodynamic radius (2.81 ±0.16 nm) derived from the cross-correlated relaxation analysis suggests that S_1_ corresponds to the TrkBtm monomer, assuming the ideal bicelle model (radius should equal 2.82 nm^29,30^) (Figure 1C,D). The S_2_ cross-peaks were observed until LPR ∼200; therefore, we can state that TrkBtm forms moderately strong dimers in DMPC/DHPC bicelles, and dimer/monomer transitions are slow in the NMR chemical shift timescale.

**Figure 1.**
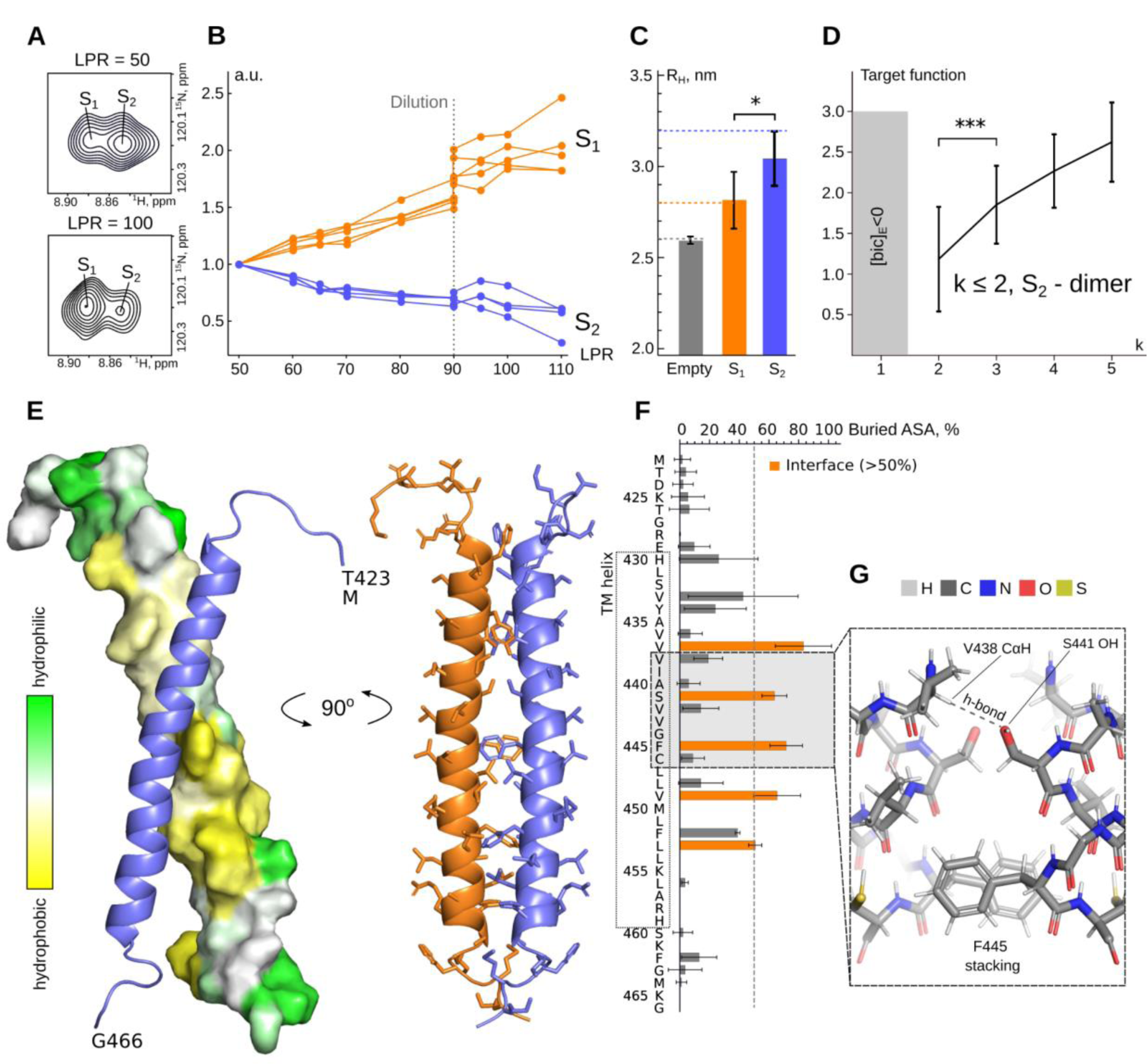
Structure of the TrkBtm dimer. **A** - The signal of the A440 amide group in the ^1^H-^15^N-HSQC spectrum of TrkBtm reveals two forms, S_1_ and S_2_, whose populations depend on the LPR. **B** - Relative intensities of the S_1_ and S_2_ backbone amide signals in the NMR spectra as a function of LPR. At LPR=90 the sample was diluted 2.5x. **C** - the hydrodynamic radii (R_H_) of an empty q=0.3 DMPC/DHPC bicelle (obtained from translational diffusion of lipids, error bar shows the Monte-Carlo SD) and S_1_ - and S_2_-containing bicelles (obtained from the rotational diffusion of proteins, error bars show the SD among the considered residues). The dashed lines denote the theoretical R_H_ values obtained using the ideal bicelle model, as described in^32^. **D** - the target function calculated for several tested oligomer orders k=2-5 of the S_2_ state^28^. In the gray area, the target function was not determined (*** indicates the statistical significance of the difference according to the t-test, p<0.001). **E** - the spatial structure of the dimeric TrkBtm. One of the helices is shown as a blue cartoon and the other - either as an orange cartoon or as a surface painted according to its hydrophobicity (using the White-Wimley scale^33^). **F** - Accessible surface area of TrkBtm residues buried in the helix dimerization interface. **G** - Closer view of the presumed key interactions supporting the TrkBtm dimer: V438 CαH-S441 Oγ polar contact and F445 stacking.

To determine the structure of the TrkBtm dimer, we placed a 1:1 mixture of ^13^C/^15^N-labeled and unlabeled proteins into bicelles with perdeuterated acyl chains at LPR 40 and used ^13^C-filtered NOESY experiments to detect protein-protein interactions directly^31^ (Figure S2). The sample containing exclusively ^13^C/^15^N-labeled TrkBtm was used as a control to exclude the effects of incomplete isotope labeling and filter leakage. This approach provided us with 13 intermolecular distance restraints, which allowed us to obtain a high-resolution structure of the protein dimer (Figure 1E-G, S3, Table 1). The protein chain forms an alpha-helix in the region H430-H459, as was already shown previously^24^ (Figure S4), and two helices engage into a parallel right-handed dimer, with a helix crossing angle equal to −33±5°, however, the helices are bent and at the place of the tightest contact this angle is equal to - 38±8°, similar to other transmembrane dimers^25^. The helix-helix interface is located at the center of the TM segment and is rather extended and nonpolar; the area of the contact surface is 493±98 A^2^. Detailed analysis revealed that dimerization is mediated by a motif with a typical 4-residue periodicity: ^437^Vxxx^441^Sxxx^445^Fxxx^449^Vxx^452^FL (Figure 1 F). Most of the residues of the motif are hydrophobic and bulky, except for S441 whose side chain seems to form a non-canonical intermolecular H-bond to the CαH moiety of V438 (Figure 1G). In addition, the F445 aromatic rings are involved in face-to-face stacking interactions.

**Table 1.** Structural statistics for an ensemble of 10 best NMR structures of the TrkBtm dimer.

| **NMR distance and dihedral restraints** |  |
| --- | --- |
| *Total NOE restraints* | 1036 |
| intra-residue | 274 |
| inter-residue | 736 |
| sequential ($\vert i - j\vert = 1$) | 278 |
| medium-range ($1 < \vert i - j\vert < 4$) | 458 |
| long-range ($\vert i - j\vert > 4$) | 0 |
| inter-monomeric NOE | 26 |
| *Hydrogen bond restraints (upper/lower)* | 156/156 |
| *Total torsion angle restraints* | 162 |
| backbone $\phi$ | 64 |
| backbone $\psi$ | 64 |
| side-chain $\chi_1$ | 34 |
| **Structure calculation statistics** |  |
| CYANA target function ($\text{\AA}^2$) | 2.12±0.38 |
| *Restraint violations* |  |
| distance ($>0.2 \text{ \AA}$) | 0 |
| dihedral ($>5^\circ$) | 0 |
| *Average pairwise r.m.s.d. ($\text{\AA}$)* |  |
| *TM helix region (430-459)<sub>2</sub>* |  |
| backbone atoms | 1.05±0.63 |
| all heavy atoms | 1.52±0.56 |
| *Ramachandran analysis* |  |
| % residues in most favored regions | 100 |
| **Helix–helix packing** |  |
| Contact surface area per dimer subunit ($\text{\AA}^2$) | 493±98 |
| Angle $\theta$ between the TMD helix axes (deg.) | -33±5 |
| Distance d between the TMD helix axes ($\text{\AA}$) | 9.20±0.45 |

### 2.1. The conformation of TrkBtm is unaffected by the environment

Since the cell membrane is mosaic^34^, and we work in the model system, it is important to test whether the parameters of the environment can affect the spatial structure of TrkBtm. First, we studied TrkBtm at a relatively low pH (6.0), below the pKa of His residues; therefore, we investigated the pH dependence of the TrkBtm NMR spectra (Figure 2A). NMR chemical shifts and the peak splitting pattern were almost identical after the pH was changed from 6.0 to physiological 7.4, suggesting that the obtained dimer structure is relevant for neutral pH.

**Figure 2.**
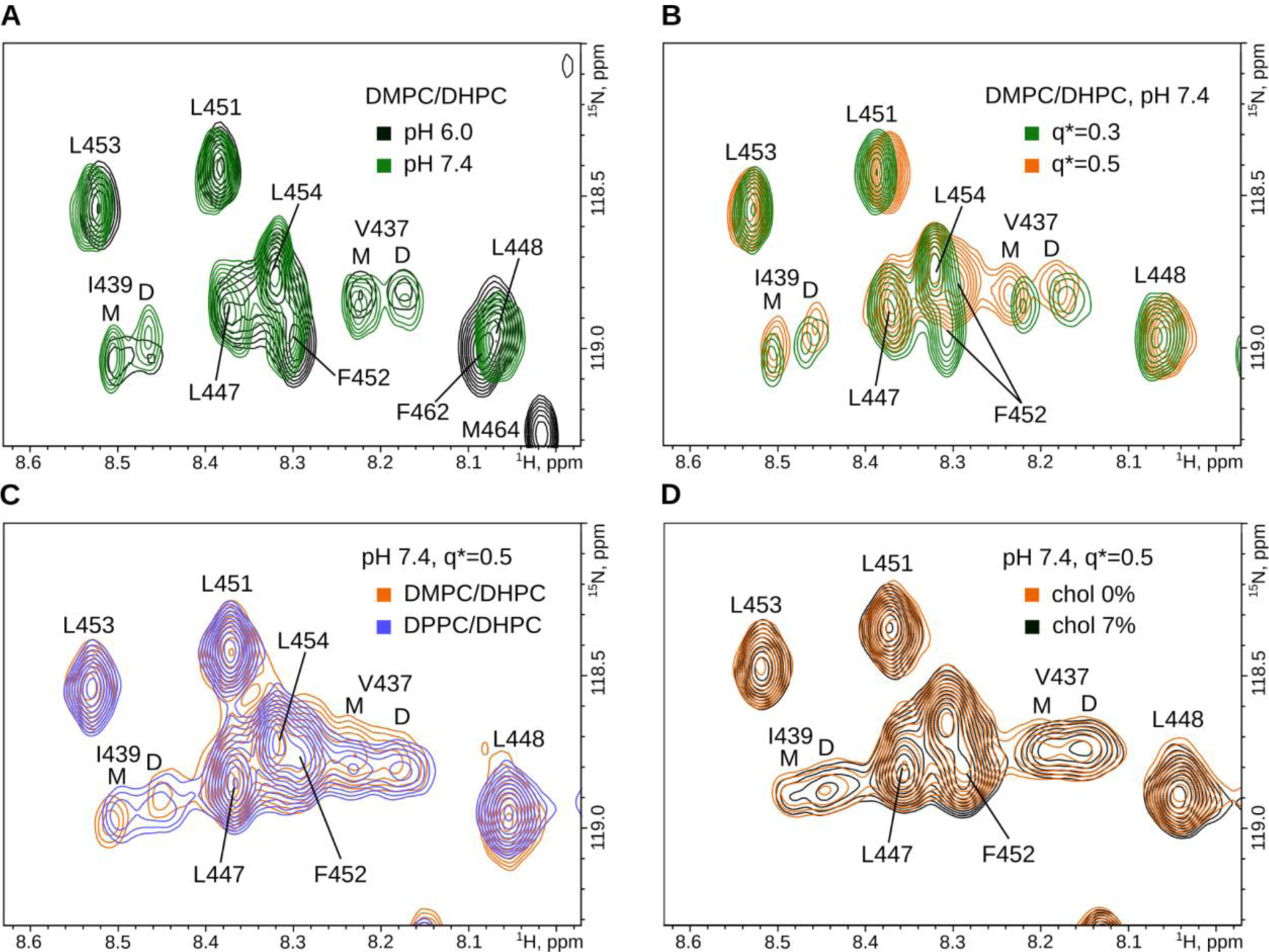
Effects of environment on TrkBtm structure. **A** - Overlaid fragments of ^1^H-^15^N-HSQC spectra of TrkBtm in DMPC/DHPC q*=0.3 bicelles at LPR=80 and pH=6.0 (black) and 7.4 (green). M and D denote the monomer and dimer signals, respectively. **B** - Spectra of TrkBtm at pH=7.4 in DMPC/DHPC bicelles formed with q*=0.3 (green) and 0.5 (orange). **C** - Spectra of TrkBtm at pH=7.4 in DMPC/DHPC (orange) and DPPC/DHPC (blue) q*=0.5 bicelles. **D** - Spectra of TrkBtm at pH=7.4 in DMPC/DHPC q*=0.5, with a bilayer containing 0 % (orange) and 7 % (black) chol (n/n).

The monomer/dimer cross-peak intensity ratio did not change as well, which means that the effect of pH on dimer stability is subtle or absent. The only noticeable change was the disappearance of signals corresponding to the unstructured terminal residue due to accelerated hydrogen-deuterium exchange.

Second, as we conducted our study in relatively small bicelles (q=0.3), we investigated whether the size of the particles was sufficient to accommodate the dimeric TM domain. Thus, we varied the bicelle q ratio in the range of q=0.3 - 0.5, which corresponds to a bicelle radius of 2.5-3.1 nm (Figure 2B). As one can see, the increase of the bicelle size has some subtle effect on the chemical shifts of certain residues, however, the pattern of peak splitting and, therefore, the dimer structure is retained.

Finally, to take into account possible variations in the bilayer content, we studied the TrkBtm in thicker bicelles, composed of DPPC/DHPC^35^ (Figure 2C). Under these conditions, cross-peaks in the NMR spectra of TrkBtm became broader because of the greater size of the particles; however, the peak splitting pattern and dimer/monomer ratio were generally the same. Thus, the found conformation of the TrkBtm dimer is highly insusceptible to the effects of the environment, which additionally supports its relevance in the case of the full-length receptor.

Another aspect of TrkB interactions with the environment is the effect of cholesterol. According to Casarotto et al.^23^, the direct addition of cholesterol to neurons may affect the BDNF-induced phosphorylation of TrkB and enhance the PLCγ1 response. This effect was dependent on the Y434 sidechain. For this reason, we titrated the monomeric TrkBtm in bicelles with cholesterol (Figure 2D, S5), introducing up to 7.0% cholesterol to the lipid vesicles, which corresponds to a 2:1 excess of cholesterol with respect to the protein. As a result, we observed subtle chemical shift perturbations (CSPs), which, however, cannot represent a high-affinity binding process because the changes are not saturated even at a 2:1 excess of cholesterol with respect to the protein. The addition of cholesterol did not influence the peak splitting pattern or population of states. On the other hand, the major CSPs were found at residues Y434 and V437, suggesting that Y434 participates in these weak interactions.

### 2.2. The found state of the TrkB transmembrane dimer is important for TrkB activation

To investigate the functional relevance of the observed binding mode, we selected several potentially critical residues and assessed their roles using site-directed mutagenesis. For the obtained structure, we selected two key amino acids at the center of the interface: S441 and F445. Additionally, we tested all TrkBtm residues with small sidechains that are known to promote helix-helix interactions: S432, A435, A440, G444, and C446. Finally, we tested the role of Y434, which has previously been shown to be important for TrkB interactions with cholesterol and antidepressant drugs^23,36^. For most of the residues, we used two types of mutations - to the residues with small and bulky side chains (Ala and Ile), to check the effect of the side chain volume (Figure 3A,B), and measured the BDNF-induced phosphorylation of TrkB. Surprisingly, only one of the tested sites substantially inhibited TrkB signaling: S441 (Figure 3D). Even an almost neutral substitution, S441A, essentially decreased the TrkB activity, and the effect of the S441I mutation was even more pronounced. This supports both the observed key position of 441 in the dimerization interface of TrkBtm and the formation of an intermolecular H-bond because the general volume and polarity of Ser and Ala sidechains do not differ significantly. In contrast, mutations of F445 to either Ala or Ile did not result in receptor inhibition, suggesting that ring stacking can be easily exchanged with tight van der Waals packing. A visible but not statistically significant decrease was also observed for S432I kinase activity. However, the position of the residue in the first turn of the TM helix suggests that changes in the TM helix structure may account for this effect. No effect was observed for the Y434 site at TrkB. Apparently, the latter residue, while being involved in the binding or sensing of lipid-like small molecules and translocation to lipid rafts, is not directly related to kinase activation and TM domain dimerization events, as previously suggested^23^.

**Figure 3.**
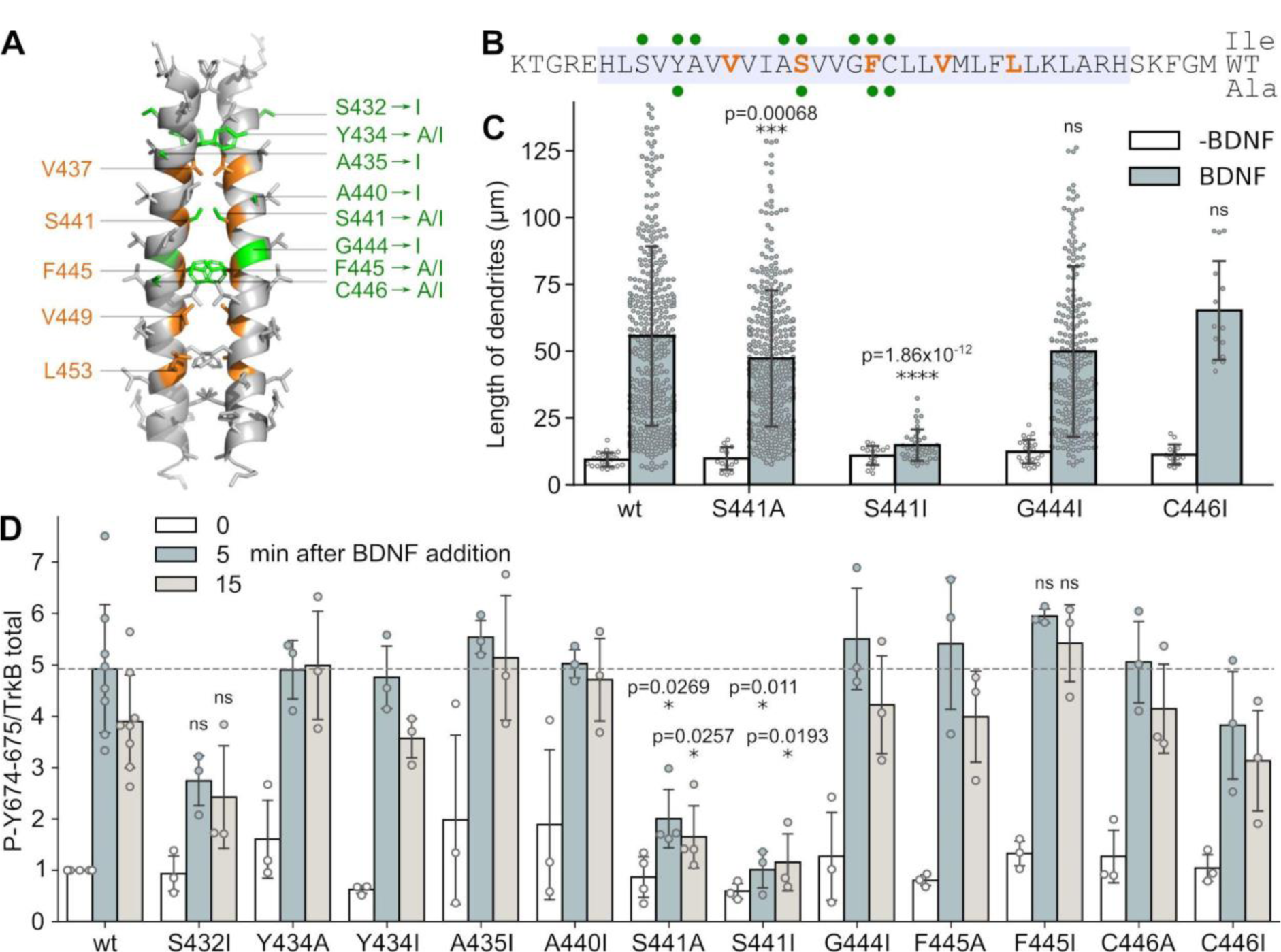
Single-point mutagenesis of TrkB transmembrane residues. **A** - Spatial structure of the TrkBtm dimer, the critical residues of the interface (orange), and the positions of point mutations (green) are indicated. **B** - Amino acid sequence of TrkBtm (highlighted in light blue), the critical residues of the interface (orange), and point mutations (green circles) to Ile (up) or Ala (down) are indicated. **C** - Average length of dendrites upon stimulation of PC12 TrkA-KO cells with BDNF. **D** - Relative amounts of phosphorylated Y674 and Y675 before and 5 and 15 min after BDNF stimulation of wild-type TrkB and its single-point mutants. Error bars denote the standard deviations, statistical significance is provided according to the Multiple t-test with Bonferroni correction (* - p<0.05, ** - p<0.01, *** - p<0.001, **** - p<0.0001, ns denotes that changes with respect to the wt are not significant).

In addition to kinase phosphorylation, we tested the direct biological activity of several selected TrkB mutants by measuring BDNF-induced cell differentiation in a specially designed cell line with the knockout of the TrkA gene^37^ (Figure 3C, S6). Both S441A and mainly S441I substitutions significantly decreased the average length of dendrites, whereas two control mutations, C446I and G444I, did not. Noteworthy, whereas the effect of S441A was statistically significant but rather subtle,/ S441I mutation was able to completely block the BDNF-induced differentiation. TrkB mutants were correctly expressed and reached the plasma membrane at similar levels as the wild type as shown by immunofluorescence of impermeabilized transfected cells and by flow-cytometry, indicating they are correctly folded and trafficked to the membrane (Figure S7, S8). Thus, we can conclude that substitutions of only those residues that are on the helix-helix interface in the reported structure provide a statistically reliable effect on TrkB kinase activity, thus validating the reported conformation as corresponding to a functionally relevant receptor state.

### 2.3. Fluoxetine interacts with the N-terminus of the TrkB transmembrane helix, stabilizing its dimeric state

Another important aspect of TrkB activity is its ability to interact with antidepressant drugs via the TM domain^23^. These properties have been studied in cells or computer simulations, but high-resolution techniques of structural biology have never been used. Here, we utilized NMR titration and chemical shift mapping to investigate the interactions between TrkBtm and drugs in lipid bicelles. We used four drugs: fluoxetine (FLX), which was previously shown to be a major drug acting on TrkB TMDs; chlorpromazine (CPR), which was shown to be inactive with respect to TrkB; and either RR or SS stereoisomers of hydroxynorketamine (RR-HNK and SS-HNK). RR-HNK was shown to bind TrkBtm, while SS-HNK was reported as a non-binder.

As a first observation, one could find out that FLX and CPR partition into the membrane mimetic (up to 92% of drugs are membrane-bound at high drug contents; at low excess of FLX, this amount is close to 100%), while HNKs do not (Figures 4 F, S9 A). In addition, both FLX and CPR induced noticeable chemical shift perturbations (CSPs) in TrkBtm in a dose-dependent manner (figure 4 A,B). However, the patterns of CSPs induced by the two drugs were different (Figure 4 C,D). While CPR induces uniform perturbations throughout the TM helix, FLX causes maximal changes at the N-terminus within the region Y434-I439. These changes were similar in both lipid environments that we tested (DMPC and DPPC bicelles, Figure S9 B). Moreover, FLX similarly affects the monomeric and dimeric states of TrkB, whereas CPR mostly affects the monomer. Notably, the highest CSPs were induced by FLX for residues Y434 and V437 in both the monomeric and dimeric states of TrkBtm, suggesting that these amino acids may be responsible for drug binding. This agrees with biochemical data that revealed the importance of tyrosine residues in FLX-induced activation of TrkB^23^. In contrast, both HNK compounds induced only subtle perturbations, with no pronounced localization. Together with the low bilayer-binding propensity of HNK (Figure 4F), this suggests that HNK could bind elsewhere and not to the TM domain, and that the effects of HNK on FLX binding to TrkB^23^ might be allosteric. Thus, we can state that FLX reveals specific interactions with the N-terminus of the TrkB TM helix in both the monomer and dimer states, while all other tested drugs do not.

**Figure 4.**
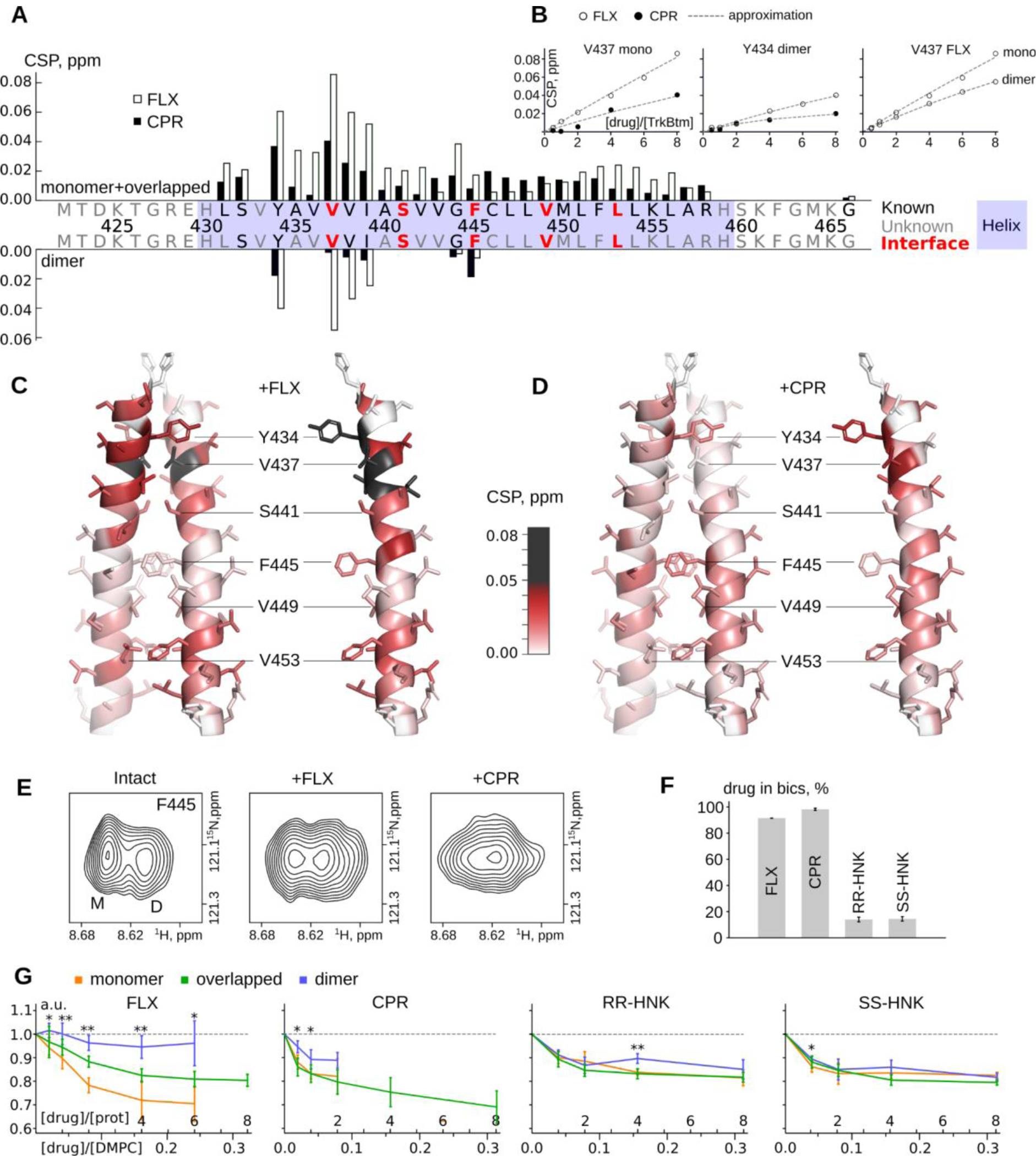
Interaction of TrkBtm with various drugs. **A** - CSPs induced in TrkBtm by FLX (empty bars) and CPR (black bars). CSPs for the separate signals of the dimer are plotted at the bottom in a mirrored scale. **B** - CSPs of V437 and Y434 plotted as a function of FLX (empty circles) and CPR (black circles) concentrations for the monomer and dimer forms of TrkBtm. The dashed lines indicate the results of a third degree polynomial approximation. **C,D-** TrkBtm dimer (left) and monomer (right) painted according to amide CSP values induced by an 8:1 excess of FLX (C) or CPR (D). Residues with CSP higher than 0.05 are shown in black. **E** - The signal of the F445 backbone amide in the intact sample (left) and with 4:1 excess of either FLX (middle) or CPR (right). **F** - Fraction of drugs residing in bicelles calculated from lateral diffusion at a drug:protein ratio of 8:1. **G** - Relative changes in the signal intensities of monomer (orange), dimer (blue), and overlapped (green) signals of TrkBtm upon drug addition. Star labels indicate statistically significant differences in the monomer and dimer populations (* p≤0.05, ** p≤0.01). The relative intensity of the monomer and dimer are plotted until the signals overlap.

Moreover, FLX stabilized the TrkBtm dimer. While the monomer intensities decreased upon the addition of FLX, the dimer cross-peaks remained unchanged, indicating an increased abundance of the dimeric state. This was not the case with all other tested drugs, which did not change the dimer/monomer intensity ratio (Figure 4G). CPR can actually destabilize the TrkBtm dimer kinetically; the addition of the compound induces the merging of two peaks corresponding to the monomer and dimer states of TrkBtm (Figure 4E, S10). This may be interpreted as an acceleration of dimer/monomer transitions, which become fast on the NMR chemical shift timescale.

## 3. Discussion

In summary, here we report the spatial structure of the TrkB dimeric transmembrane domain. The obtained structural data need to be discussed in the context of current information present for Trk receptors, which includes the structure of the TrkA transmembrane domain^20^ and model of TrkB TMD, obtained by computer simulations^23^. Notably, the aforementioned model is quite different from the spatial structure reported here. In that model, the dimer is formed via the canonical “glycophorin” GxxxG-like motif^38^ (^440^Axxx^444^G), which is supported by Y434 at certain cholesterol contents, and the dimer topology is left-handed. According to our data, the dimer is formed via the extended motif, with the key residue being S441, and with a right-handed mode of helix-helix interaction. The latter conformation was supported by functional assays; the S441A and S441I mutations drastically inhibited the BDNF-induced activation of TrkB, while the A440I and G440I substitutions did not demonstrate any pronounced biological effect. Therefore, we conclude that the observed structure of the TrkBtm dimer most likely corresponds to the active state of the receptor. The alternative conformation, proposed by computer modeling, may in turn describe the dimeric but inactive state, which should also be a stage of TrkB signaling, according to the most recent findings^18^. The left-handed state may be also be favored by the high cholesterol contents in the membrane, in agreement with the in silico data, which explains the inhibiting effects of high cholesterol excess on TrkB.

To further understand the transmembrane domain conformations that occur in the course of neurotrophin signaling, one could invoke data regarding TrkA, one of the closest TrkB relatives. The spatial structure of its TM domain most likely represents the inactive dimer state, as shown by cross-linking^20^ and further by NMR experiments^11^. The active state of TrkA TMDs can be obtained by cross-linking at the K410 position, but its exact conformation has not been reported. Comparing the two structures one could find out that they are drastically different (Figure 5). First, the TM helix of TrkB is substantially (six residues) longer (Figures 5B,C, S11), which questions the appropriateness of such a comparison. This difference is not an artifact of membrane mimetics, because the length of the helix in TrkA does not depend on the choice of the environment and is equal in both micelles and bicelles^11,20,30^. It seems that while the structures of the ectodomains in complex with their ligands are almost identical ^7,8^, the transmembrane parts of these two related proteins diverged strongly during evolution. The sequence comparison of TMDs revealed almost no identity when looking at the distribution of small and aromatic residues that can promote dimerization, while a high identity level was found between the TM regions of TrkA and TrkC (Figure 5C). This statement agrees with the results of functional studies. TrkA and TrkC were found to cause cell death, but the TrkB receptor was not, and the key role in this effect was played by the TM domain^39,40^. Thus, the parameters of transmembrane states may be not uniform within the Trk family.

**Figure 5.**
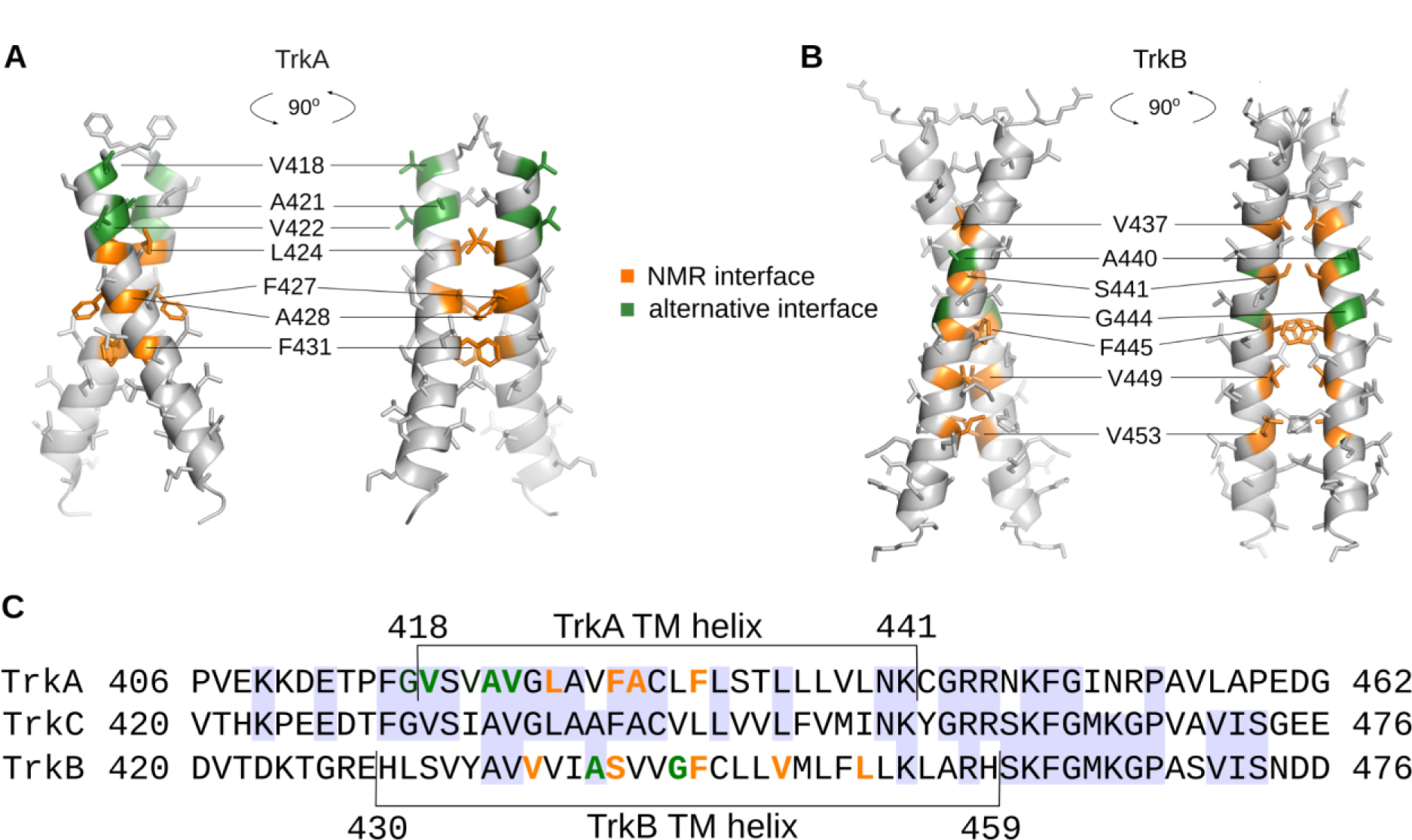
Comparison of TM domain dimer conformations within the Trk family. **A,B** - the spatial structures of TrkA (A, PDB 2N90) and TrkB (B, this work) TM dimers obtained by NMR in DPC micelles and DMPC/DHPC bicelles, respectively. Orange and green colors denote the residues on the experimentally determined interface and on the interface, assumed from the cross-linking experiments (A) or computer modeling (B). **C** - Manual sequence alignment of Trk family members based on the Clustal Omega program (http://www.clustal.org/) results for alignment of the transmembrane and intracellular domains. Experimentally determined TM domains are indicated, blue background highlights the identical residues.

Next, one can find that the structures of the transmembrane domains are also essentially distinct. In the case of TrkA, the dimer is left-handed, supported by stacking interactions and van der Waals packing, while the TrkB dimer is right-handed and supported by a polar contact of Ser sidechain. This difference may be ascribed to the different types of membrane mimetics that were used to accommodate the protein, and the ability of a dimeric TM domain conformation to switch between two states when the protein is studied in micelles and in bicelles ^25–27^. Interestingly, the TrkBtm dimer structure obtained by computer simulations is quite similar to the TrkA structure (left-handed mutual arrangement of TM segments), supporting our hypothesis that the state could correspond to the inactive pre-formed dimer of TrkB. In this regard, one could assume that if the TM domain conformations are uniform within the family, signaling is accomplished by switching between the left-handed and right-handed conformations of Trk TM domains, which is accompanied by mutual rotations of the TM helices.

Finally, we analyzed the interactions between TrkBtm and antidepressant drugs as well as cholesterol. The effects of cholesterol are unclear. In our model system, we did not observe any significant events that could be interpreted as strong and specific interactions between the protein and cholesterol regardless of the presence of the CRAC motif^41^. The maximal chemical shift perturbations were twice lower than those induced by FLX at the same ligand/protein and ligand/lipid ratio but were comparable to the ones induced by the CPR (Figure S9 C,E). Unlike FLX, cholesterol stabilized neither the monomer nor dimer of TrkBtm (Figure S9 D). Additionally, the changes were not saturated even under excess cholesterol. This is in agreement with the computer modeling data which suggests that the effects of cholesterol on TrkB are mostly due to variations in bilayer thickness^23^. However, weak interactions involving the Y434 sidechain were indeed detected. Taking into account the high overall cholesterol contents in membranes (30%), these weak contacts may become relevant at physiological cholesterol concentrations. Our data indicate that the effects of cholesterol on TrkB signaling are accomplished not via the highly specific interactions of cholesterol with the CRAC motif, but most likely by the effect of cholesterol on the physical properties of the cell membrane, its ability to maintain the liquid-ordered state of the bilayer and regulate the abundance of lipid microdomains. The partitioning of proteins between domains may be modulated by the CRAC domain, which explains both the effects of cholesterol on cells and the effect of the Y434 side chain. This also means that the mere presence of the CRAC/CARC motif does not ensure specific protein-cholesterol interactions. Alternatively, the CRAC motifs could become important when TrkB enters the lipid rafts of the cell membrane upon the BDNF binding^42^, and the weak interactions that were found here may become strong within the liquid-ordered bilayer. Unfortunately, it seems hardly possible to mimic this state within lipid bicelles, and computer modeling is unable to reproduce it in systems with a low number of lipid molecules, leaving the question open for future studies.

As for drug interactions, our spatial structure helps us to understand the structural basis of their activity. Here, we show that FLX interacts with the N-terminus of the TrkB TM helix in both the dimer and monomer states, supporting all previous data obtained on cells^23^. The interaction does not change the structure of the TrkBtm dimer but rather stabilizes this state compared to the monomer. These features are somewhat different from the previously suggested mechanism, stating that FLX binds to the dimeric state of TrkB and alters the distance between the C-termini of the TMD helices^23^. However, these two hypotheses may be combined to provide a self-consistent view of the FLX mechanism, assuming that the AxxxG conformation of TrkB corresponds to the receptor inactive state. According to both modeling and NMR, the site of FLX binding to TrkB involves residues Y434 and V437, which interfere with the structure of the AxxxG TrkB dimer (inactive state). At the same time, FLX binding to TrkB does not conflict with the active dimeric state reported here, and, moreover, stabilizes it. Altogether, this would mean that FLX disturbs the formation of an inactive conformation and supports the formation of an active dimeric arrangement of TMDs, which agrees perfectly with the observed biological effects of FLX - the activation of the TrkB receptor. None of these features of FLX were observed for the antipsychotic CPR, which correlates with the absence of antidepressant effects. Besides, the absence of Y434 or similar residue in TrkA, explains why the activity of this protein is not sensitive to FLX.

To summarize, we report here the spatial structure of the dimeric transmembrane domain of TrkB in lipid bicelles. The obtained conformation is stable in a variety of environments and corresponds to the receptor’s active state, according to the directed mutagenesis. Together with previously published results of computer simulations, our data helps to build a self-consistent model, explaining the activity of FLX with respect to the TrkB receptor.

## Acknowledgments

The work was supported by the Russian Science Foundation grant #22-14-00130 to SAG (NMR studies), to the Spanish Ministery of Science and Innovation grant number PID2021/127600NB-I00 to MV. ABM is the recipient of an Investigo contract from the Generalitat Valenciana (INVEST/2022/456) and by EU funding within the NextGeneration EU-MUR PNRR TUSCANY HEALTH Ecosystem – THE (Project no. ECS_00000017)’ spoke 1 and 8 to AC. We are thankful to Prof. Eero Castren for the fruitful discussion of the manuscript.

## 4. Methods

### 4.1. Protein sample preparation

The details of protein synthesis, purification, and sample preparation for NMR analysis are provided in our most recent work^24^. The gene encoding the transmembrane domain of the human TrkB (M^423^TDKTGREHLSVYAVVVIASVVGFCLLVMLFLLKLARHSKFGMK^466^G UNIPROT Q16620), was synthesized by Cloning Facility (Russia) with codon optimization for *E. coli* and cloned into pGEMEX-1 vector using NdeI and HindII restriction sites. The protein was synthesized by a cell-free continuous exchange expression system. A standard cell-free reaction (2-3 ml of RM) was carried out in 50 ml tubes using a Pur-A-Lyzer Maxi dialysis kit (#PURX35050, Sigma). The TrkBtm was expressed as a precipitate. The algal mixture of ^15^N (#NLM-6695, CIL) or ^13^C-^15^N-labeled (#CCN070P1, CortecNet) amino acids and ^15^N (CN501P05, CortecNet) or ^13^C-^15^N-labeled (CCN500P1, CortecNet) cysteine were used to obtain a uniformly ^15^N or ^13^C-^15^N-labeled protein samples, respectively. The precipitate from the cell-free reaction was washed 3 times with water (milliQ), solubilized with detergent buffer (20 mM Tris 8.0, 150 mM NaCl, 10 mM bME, 1 mM EDTA and 1.5% lauryl sarcosine), purified using size-exclusion chromatography in SEC-buffer (20 mM Tris 8.0, 50 mM NaCl, 5 mM bME, 0.5% lauryl sarcosine) and target protein was precipitated by a TCA/acetone procedure^43^. The dry powder of ^15^N or ^13^C-^15^N-labeled TrkBtm was dissolved in 450 μl of trifluoroethanol(TFE)-water mixture containing 45 μl deuterated TFE for frequency lock, 1mM of TSP-d4 as a reference compound and 1 mM Tris(2-carboxyethyl)phosphine to prevent disulfide bonds between the transmembrane C451 residues. As TrkBtm contains no Trp residues and the optical concentration measurement is impossible, 1D ^1^H NMR spectra were acquired to estimate the concentration of protein using signals of the TSP methyl group at 0.0 ppm as a reference. Then, we solubilized TrkBtm in bicelles as described in our recent work^44^. Briefly, deuterated or protonated DMPC, DPPC, and DHPC were added in needed amounts to the sample, and water was added up to a TFE/water ratio of 1:1. Then the sample was freeze-dried for 12-15 hours and dissolved in 450 or 330 μl 20 mM phosphate buffer pH 6.0 or 7.4 containing 5% D_2_O for frequency lock, 0.01% NaN_3_, 0.5 mM TSP-d4 as a chemical shift reference and 2 mM EDTA.

### 4.2 NMR spectroscopy and spatial structure calculation

NMR spectra were acquired at 40°C on Bruker Avance III 600 MHz and 800 MHz spectrometers equipped with triple resonance cryogenic probes. The ^13^C-^15^N-labeled samples (330 ul) for structure calculations were placed into a 5-mm Shigemi tube to maximize the protein concentration. The rest of the samples (450 ul) were placed into regular 5-mm glass tubes (Wilmad, USA).

To assign the ^1^H, ^13^C, and ^15^N resonances and resolve the structure of a TrkBtm monomer we recorded the set of 2D heteronuclear (^1^H-^15^N-HSQC, ^1^H-^13^C-HSQC aliphatic and ^1^H-^13^C-HSQC aromatic), 3D triple-resonance NMR spectra (HNCA, HNCO, HNcoCA, HNcaCO, HNCACB, NOESY-HSQC), 3D ^13^C- and ^15^N-NOESY-HSQC and 3D-HccH-TOCSY spectra. Aromatic sidechains were assigned based on the 2D hC(CC)H and 3D (H)CCH-COSY experiments^45,46^. Spectra were recorded, where possible, using the BEST-TROSY pulse schemes^47^ and non-uniform sampling in indirect dimensions. Such spectra were processed with qMDD using the “iterative soft thresholding” algorithm with virtual echo^48^.

The spatial structure of the TrkBtm dimer was calculated using the CYANA program^49^. Backbone torsion angle restraints were obtained based on the NMR chemical shifts in TALOS-N software^50^. Sidechain angle restraints were found by manual analysis of NOESY spectra and ^3^J_CGCO_ and ^3^J_CGN_ vicinal J-couplings that were measured using the spin-echo difference constant-time HSQC NMR experiments^51,52^. To obtain the intermolecular distance restraints, we recorded two 3D ^13^C-filtered-NOESY-HSQC spectra for the samples containing the 13C/15N-labeled TrkBtm and an equimolar mixture of isotope-labeled and unlabeled proteins (Figure S2).

To measure the translational diffusion of DMPC, DHPC, and drugs we employed the ^1^H pulsed-gradient stimulated echo pulse sequence with convection compensation^53^. In this experiment, 16 1D spectra were recorded with the gradient power increasing from 13.9 to 53.0 Gs/cm. We fitted the peak intensity decays using the Wolfram Mathematica 5.0 software and obtained the diffusion coefficients to calculate the hydrodynamic radii as described in our previous work^54^.

The hydrodynamic radii of TrkBtm in bicelles we calculated from the rotational correlation time τc:

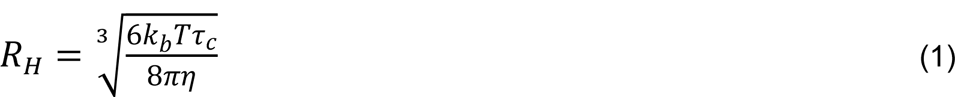

Here, kb is the Boltzmann constant, T is the absolute temperature and η is the dynamic viscosity of the solution. The τ_c_ values were derived from the cross-correlated relaxation rate as described elsewhere^55^ assuming that ps-ns range internal motions were negligible in the helical TrkBtm backbone. Cross-correlation relaxation rates were measured at LPR=60.

Obtained structures were analyzed using MOLMOL^56^ and PyMOL (Schrödinger LLC) software.

### 4.3. Oligomer order determination

In the current work we used an NMR-based approach to describe complex oligomer equilibria of TMDs described earlier^28,32^. Briefly, we applied the so-called “micellar solvent” model (which can be applied to bicelles as well)^57^. According to this model, the equilibrium constant K_eq_ of n monomers association into an n-mer depends on the concentrations of monomer [M], n-mer [N] and empty bicelles [bicE] in the following way:

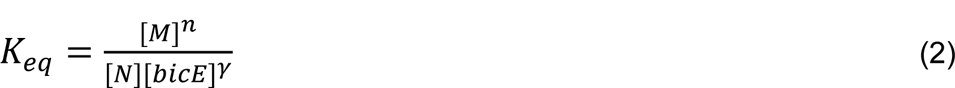

Here γ is the effective order of reaction with respect to the concentration of empty bicelles. Thus, if we sequentially add bicelles to the sample, determine [M] and [N]/n according to the intensities, and plot [M]^N^/[N] as a function of [bicE] it would be a straight line in logarithmic coordinates with the tilt γ. If we then dilute the sample and continue adding bicelles, the points after dilution would fit the same line only if the oligomer order n was assumed correctly^57^.Thus, to determine the oligomer order we used a target function that measures the deviations of points after the dilution from the initial dependence^28^. This target function was determined for several peaks of TrkB assuming different oligomer orders and the distributions were compared using the Student t-test.

Besides, we took into account the size of the bicelles, which was measured from the translational diffusion coefficients of lipids for the empty bicelles and from the rotational diffusion coefficients that were obtained based on the protein ^1^H,^15^N cross-correlated relaxation rates for the bicelles bearing the oligomers of TrkBtm as described^30^. Diffusion coefficients were then translated into the hydrodynamic radii using the Stock-Einstein relationships and then compared to the values predicted based on the “ideal bicelle model”, assuming that the bilayer thickness is 4.0 nm^58^ and bicelle surface area that is occupied by one TM helix is 1.4 nm^2^ ^30^.

### 4.4. TrkB interactions with small molecules

We performed a gradual titration of TrkBtm with 10 mM FLX and 40 mM CPR, RR- and SS-HNK solutions in a 20 mM phosphate buffer pH 7.4. These solutions contained DHPC at a concentration of 8.4 mM to keep the bicelle q* (effective lipid/detergent ratio taking into account that some detergent is present in the monomeric state) and size constant upon titration^30^. As CPR was insoluble at this pH, it was added as a well-shaken emulsion which turned transparent and solubilized well in the bicelles. The amount of added CPR was controlled by ^1^H NMR spectra.

To observe the partitioning of drugs between the solution and bicelles with TrkBtm we measured the diffusion coefficient of DMPC (D_DMPC_) and drugs (D_drug_). In a separate experiment, we measured the diffusion of 1 mM of each drug in a buffer containing 8.4 mM DHPC, which corresponds to the concentration of monomeric DHPC in the presence of q=0.5 DMPC/DHPC bicelles(D_drug,free_)^30^. Assuming that D_DMPC_ characterizes the whole bicelle, D_drug_ depends on the distribution between bicelles and solution the following way:

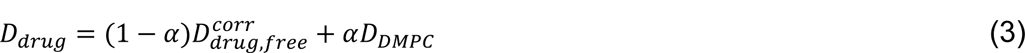

Here α is a share of drug in bicelles and D_drug,free_ is corrected to include the effect of volume fraction Ф occupied by bicelles^59^:

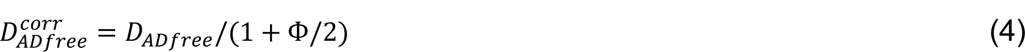

Finally, we can calculate α:

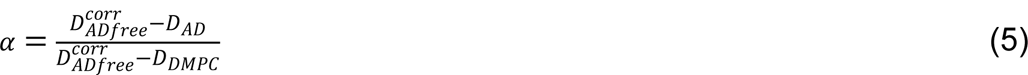

To characterize the interactions between the small molecules and TrkBtm, we measured the CSPs and populations of monomers and dimers. CSPs were calculated in the following way:

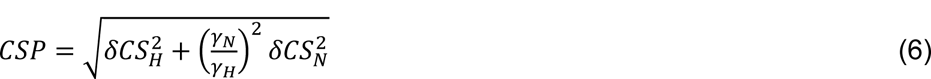

Here δCS_H_ and δCS_N_ are the changes of ^1^H and ^15^N chemical shifts, γ_H_ and γ_N_ are the gyromagnetic ratios of ^1^H and ^15^N nuclei, respectively.

To observe the changes in monomer and oligomer populations, we measured the intensities of the separately standing cross-peaks of TrkBtm, corresponding to the monomeric and dimeric states in ^1^H,^15^N-HSQC spectra. All the intensities were normalized to the values measured in the absence of the drug and to the number of scans in each spectrum. The values for each drug ratio were averaged and the SD values were calculated.

Cholesterol was added to the sample as an 85 mM solution in ethanol (33 mg/ml) at 45 °C. We added cholesterol three times, monitoring its incorporation into the bicelles by following the signals of its methyl groups in the ^1^H spectra (Figure S4 A). At each titration point some cholesterol precipitated, and at the 3rd point, no cholesterol was further retained in the solution. To measure the exact contents of cholesterol in bicelles we redissolved the supernatant of the sample after chol titration in CDCl_3_:CD_3_OD 1:1 with the addition of 1mM TSP-d4 and applied the quantitative NMR.

To distinguish between the effects of cholesterol and ethanol we performed a separate experiment preceding the cholesterol titration. We reproduced the ethanol contents (2% v/v) at the last point of cholesterol titration and measured the CSP and intensity alterations. The ethanol-free CSP induced by cholesterol was calculated as a length of the vector difference between C_2_D_5_OH and C_2_D_5_OH+chol effects as follows:

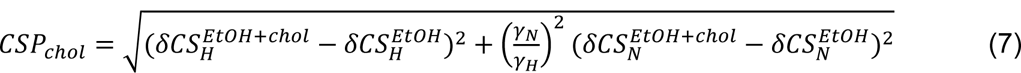

### 4.5. DNA constructs for functional assays

A plasmid encoding rat TrkB with an N-terminal hemagglutinin tag was kindly provided by Dr. Y. Barde. All TrkB mutants and constructs were derived from this plasmid. Mutagenesis was done using the site-directed mutagenesis kit (Agilent) according to the manufacturer’s protocol. The oligonucleotide sequences of all of the constructs are available upon request. All DNA constructs were sequenced at Macrogen.

### 4.6 Cell culture and transfection

HeLa cells, which do not express endogenous TrkB, were cultured in Dulbecco’s modified Eagle’s medium (Fisher) supplemented with 10% fetal bovine serum (Fisher), 1% Penicilin/Streptomicin and 1% L-Glutamine at 37 °C in a humidified atmosphere with 5% CO2. PC12-TrkA-KO (REF) cells were cultured in Dulbecco’s modified Eagle’s medium with 10% fetal bovine serum, 1% Penicilin/Streptomicin and 1% L-Glutamine and 5% horse serum (Fisher). Transfection of cells was performed using Lipofectmine-2000 following the manufacturer’s protocol.

### 4.7 Cell signaling

HeLa cells were seeded in 60 mm dishes and transfected with 3 ng of DNA. Twenty-four hours after transfection, the cells were lifted and replated into 12-well plates at a density of 100,000 cells/well. Forty-eight hours after transfection, the cells were starved in serum-free medium for 2 h and were then stimulated with 50 ng/ml BDNF (Alomone) at the indicated time intervals. The cells were lysed with TNE buffer (Tris-HCl, pH 7.5, 150 mM NaCl, 1 mM EDTA) supplemented with 1% Triton X-100 (Sigma), protease inhibitors (Roche), 1 mM phenylmethylsulfonyl fluoride (Sigma), 1 mM sodium orthovanadate (Sigma), and 1 mM sodium fluoride (Sigma). Lysates were kept on ice for 10 min and centrifuged at 12,000 × g for 15 min in a tabletop centrifuge. The protein level of the lysates was quantified using a Bradford kit (Pierce), and lysates were analyzed by SDS-PAGE. The proteins were resolved in SDS-PAGE gels and transferred to nitrocellulose membranes that were incubated overnight at 4 °C with one of the following antibodies: mouse monoclonal anti-hemagglutinin (1:2000, Sigma); rabbit anti-P-TrkB-Tyr^674/675^ (1:1000, Cell Signaling). Following incubation with the appropriate secondary antibody, anti-mouse (1:10000, Invitrogen), and anti-rabbit (1:10000, Invitrogen), the membranes were imaged using LICOR. Protein bands were quantified by the LOCOR lite software.

### 4.8 Differentiation of PC12-TrkA-KO cells

The generation of the PC12-TrkA-KO cells was described in^37^. Transfection in PC12-TrkA-KO cells was performed using Lipofectamine-2000 as previously described. Cells were plated into 24-well plates and transfected with 1µg of plasmids encoding TrkB or TrkB mutants and 0,2 µg of GFP. Transfection was allowed for 24 hours and cells were washed three times with serum-free medium and incubated for 48 h in a medium containing 1% fetal bovine serum and 20 ng/ml of BDNF (Alomone). At 48 h, the cells were washed with cold PBS and fixed with 4% paraformaldehyde for 15 min at room temperature. DAPI was used for nuclei staining and samples were mounted using mowiol medium. The cells were imaged using a Leica SP8 spectral confocal microscope. The length of the longest neurite was measured using Image J, data were analyzed and plotted with GraphPad Prism 8.

### 4.9 Cytometry and immunofluorescence

Membrane localization of TrkB mutants in Hela cells was analyzed by Flow cytometry and by immunofluorescence in non-permeabilized cells. Hela cells were transiently transfected with the indicated TrkB and GFP constructs. 48 h later, the cells were fixed with 4% paraformaldehyde, blocked with 10% FBS for an hour, and incubated with anti-HA antibody (1:200, Sigma, H9658) overnight. The following day, cells were washed with PBS and incubated with the anti-mouse (1:500, Invitrogen, A31570) for an hour. Nuclei were stained with DAPI and imaged using SP8 Leica confocal microscope. Alternatively, membrane localization of TrkB mutants in Hela cells was analyzed by Flow Cytometry. Briefly, 48h after transfection, Hela cells were lifted and incubated with PBS with an anti-HA antibody (1:1000, Sigma, H9658) for an hour, followed by incubation with a secondary antibody labeled with Alexa-647 (1:100, Invitrogen, A21447). GFP-positive cells were selected and the Alexa-647-positive cells were quantified using the Flow Cytometer MACS Quant 10 (Miltenyi).

## Data availability

Spatial structure and chemical shifts were deposited to BMRB under the access code 34814 and to PDB, ID 8OYD.

## Author contributions

E.F.K. conducted the NMR experiments and analyzed the data, S.A.G. synthesized the protein and analyzed the data, M.L.F. and A.B.M. constructed the TrkA mutants and performed the functional assays, M.C. and A.C. created the knockout cell line, K.S.M, A.S.A., S.A.G. and M.V. designed and supervised the project, K.S.M. wrote the paper with the assistance from all the authors, S.A.G., A.B.M, M.V. and A.C. acquired the funding.

